# Identification of 5mC within heterogenous tissue using *de-novo* somatic mutations

**DOI:** 10.1101/2024.06.05.597613

**Authors:** Justin Jon Schader Wilcox, Quentin Jean Robert Foucault, Toni Ingolf Gossmann

## Abstract

Tissues represent a fundamental evolutionary interface at the junction of genotype and phenotype. Indeed, gene regulation often occurs at the tissue level and manifests itself through tissue-specific epigenetic modifications. Studies investigating tissue epigenetics are limited by access to pure tissues. Tissues not only differ epigenetically, they are also subject to genetic differentiation through somatic mutations. As somatic mutations follow predictable patterns of inheritance, the application of population genomic approaches to inter- and intra-tissue variation could allow for the efficient detection of epigenetic modifications, even when tissue samples are convoluted. Here, we present an approach that uses *de-novo* somatic mutations to deconvolute 5mC methylation patterns through analysis of shifts in tissue-specific allele frequencies. We use simulations and bisulfite sequencing data to show that somatic mutations are common and detectable in next-generation sequencing data. We then use changes in mutation frequencies to accurately derive the proportional tissue of origin along a gradient of *in silico* subsamples of mixed-tissue bisulfite reads. We confirm that mixed tissues bias estimates of methylation levels and prevent detection of methylation differences at high levels of mixture. Our derived estimates of tissue contamination allow for unbiased and accurate deconvolution of mixed-tissue methylations in CpG and non-CpG context. We are ultimately able to recover 15-30% of differentially-methylated sites, and approximately 40-90% of differentially-methylated CpG islands and gene bodies in any cytosine context at contamination levels up to 90%. Our findings highlight the utility of population genomic approaches across scales, and expand the accessibility of epigenetics studies within evolutionary biology.

## Introduction

The tissues of multicellular organisms constituent a fundamental and neglected evolutionary interface at the junction of phenotype and genotype (Barbeira et al., 2018; Cardoso-Moreira et al., 2019; Dalziel et al., 2009). From a theoretical perspective, selection acts on phenotypic differences, which results in part from differences in gene expression (El Taher et al., 2021; Hill et al., 2021). Gene expression is fundamentally regulated at the level of cell-types and tissues (Kim-Hellmuth et al., 2020; Kryuchkova-Mostacci & Robinson-Rechavi, 2017; Sonawane et al., 2017), making questions of cellular identity and tissue-differentiation central to the study evolution in multicellular life (Banta & Richards, 2018; Blake et al., 2020; Kryuchkova-Mostacci & Robinson-Rechavi, 2015). Studies on gene expression and evolution have, however, historically focused on whole organisms (Ashe et al., 2021; Herrel et al., 2020; Laine et al., 2023; Signor & Nuzhdin, 2018), divorced from mechanistic questions of tissue differentiation, physiology and development. Evolutionary studies incorporating tissue-specific effects are therefore of emerging interest (Oliva et al., 2020; Sebe-Pedros et al., 2018; Tanay & Sebé-Pedrós, 2021; Tosches et al., 2018).

Epochal advances in genomics now provide an opportunity to expand evolutionary studies across the scales of organisms, tissues and development (Marioni & Arendt, 2017; Svensson et al., 2021; Tickle & Urrutia, 2017). Tissues are composed of distinct cell lineages (Woodworth et al., 2017), which are subject to their own somatic evolution (Lupski, 2013; Oota, 2020). Advances in sequencing and cell-imaging technology have made it possible to identify and track the genomic changes that occur during tissue differentiation (De, 2011; Fernandez et al., 2016; Lupski, 2013; Vattathil & Scheet, 2016). Mosaic changes in DNA sequences are persistently observed in the tissues of multicellular organisms, and include somatic mutations (Cagan et al., 2022), structural variants (Erickson et al., 2022; Yu et al., 2020), transpositions (Loreto & Pereira, 2017), and even tissues-specific karyotypes (Kinsella et al., 2019; Rezaei et al., 2020). Although long recognized as important in cancer biology, the potential biological significance of such changes has yet to be fully evaluated.

Distinctions between tissues are more traditionally understood from the perspective of epigenetics: non-mutational DNA modifications, which give rise to differences in the expression of RNA and ultimately translation of proteins (Atlasi & Stunnenberg, 2017; Tanay & Sebé-Pedrós, 2021). Though varied in form, such modifications to either nucleotide bases and/or DNA packaging (Carter & Zhao, 2021; Paksa & Rajagopal, 2017) provide an essential regulatory framework for differential gene expression between heterogeneous cells and tissues. Indeed, in humans, these modifications allow clear diagnostic identification of tissue types, origins, and pathologies (Loyfer et al., 2023; Moss et al., 2018). Though generally conserved in cell lineages, patterns of epigenetic modification and inheritance are highly variable and unpredictable among eukaryotes even at relatively fine taxonomic scales. As such, epigenetic models for the identification of cell-lineage and tissue origin are only possible in well-studied, primarily model, taxa (Tanay & Sebé-Pedrós, 2021). Characterization of epigenetic modifications and their roles in development of non-model organisms remain important to understanding acclimation, adaptation, and comparative evolution (Lamka et al., 2022; Sadler, 2023).

Efforts in applied and comparative epigenetics have largely revolved around CpG sites and 5mC modification. Though not universal, methylation of cytosine to form 5-methyl-cytosine (5mC) is among the best-studied forms of epigenetic modification (de Mendoza et al., 2020), and has formed a focal point for the study of gene regulation in model and non-model organisms alike (Sadler, 2023; Schmitz et al., 2019). They occur in at least four of the five supergroups of Eukaryotes—SAR (Hoguin et al., 2023), Archaeplastida (Zhang et al., 2018), Amoebozoa (Fisher et al., 2004), and Opisthokonta (Bewick et al., 2019)—but are poorly conserved within these (Aliaga et al., 2019). For example, they are common in deuterostomes but absent from many protostome lineages (de Mendoza et al., 2020). In vertebrates 5mC modifications are associated with repression of gene expression and are most pronounced in CG dinucleotide (CpG) context (Schmitz et al., 2019). These mechanisms complement the effect of regions of densely grouped CpG sites—known as CpG Islands (iCpG)—to enhance transcription, as part of a unified and vital system to control gene expression in many organisms (Angeloni & Bogdanovic, 2021). This system can, however, be unpredictable even within well-studied amniotes (Luo et al., 2018): 5mC can commonly occur in non-CpG context and have incongruous impacts across genomic regions, tissues, and organisms (Derks et al., 2016; Fuso, 2018; Ramasamy et al., 2021). As such, further research is needed on the diversity, the function and role of 5mC methylation, particularly in non-model organisms.

Studies on 5mC methylation are however constrained by two factors: difficulties in the isolation of pure tissues (Lamka et al., 2022; Luo et al., 2018; Sturm et al., 2021), and incomplete knowledge of the occurrence and distribution of 5mC across eukaryotic genomes (Tanay & Sebé-Pedrós, 2021). Highly vascularized tissues, such as those from kidneys, spleen, and adipose can often be contaminated with blood (Krausgruber et al., 2020; Liu et al., 2023; Vogel et al., 2019). Fetal, embryonic and germline tissues (Eckmann-Scholz et al., 2012; Jenkins et al., 2017; Morin et al., 2017)—particularly important to understanding tissue development (Ashe et al., 2021)—and tumors—for which breakdowns in epigenetic regulation are common—pose even greater challenges (Bergmann et al., 2016; Taylor-Weiner et al., 2018).

Deconvolution approaches have been applied to address cross-tissue contamination, but these are generally reliant on prior species-specific knowledge of 5mC methylation patterns: Reference-based deconvolution using atlases of well-documented epigenetic modifications by cell type are limited to very-well-studied organisms—primarily humans (Loyfer et al., 2023; Moss et al., 2018; Tanay & Sebé-Pedrós, 2021); multivariate ordination-based deconvolution techniques rely on multiple samples and make assumptions of consistent modifications in the tissues of these samples (Houseman et al., 2016; Scherer et al., 2020); and single-sample techniques are restricted to assumptions of consistent CpG methylations in the context of CpG islands (Jeong et al., 2022; Zheng et al., 2014). None of these methods are appropriate to the deconvolution of tissues in non-model organisms in which inconsistent epigenetic inheritance and non-canonical patterns of cytosine methylation may occur. Analyses of mosaic genomic heterogeneities based on the application of generalized evolutionary principles could, however, allow for nearly-universal deconvolution of tissues in any organism.

The genomic resources available for the great tit, a songbird widespread in Eurasia, include a high-quality reference genome as well as whole genome bisulfite data from the same individual. This data set provides a superb system for studying both the detection of somatic mutations and their use in deconvolution. The data set contains, bisulfite reads from two tissues—blood and brain—as well as deep sequencing of blood by unmodified Illumina sequencing, allowing for independent validation of somatic mutation detection. As all data are from the individual that was used to assemble the reference genome, all single nucleotide variants (SNVs) should be either germline-heterozygous or somatic mutations in all data types. Finally, great tits are known to have non-canonical methylations of cytosine in non-CpG context, and (unlike mammals) poorly retained patterns of CpG methylation across cell divisions (Derks et al., 2016), making them emblematic of the sorts of non-model organisms ineligible for deconvolution with existing applications.

Here we propose a method for the deconvolution of paired-tissues using allele frequencies of somatic mutations and apply it to 5mC data. We first document the feasibility of using somatic mutations to deconvolute mixed-tissues by simulating allele frequencies for somatic mutations and contrast them to true germline heterozygous sites. After demonstrating the capacity and expected distributions of somatic mutations using simulations, we then document the convolution problem and validate our proposed solution using a gradient of *in silico* mixtures of blood and brain whole-genome bisulfite (WGBS) sequencing data taken from the great tit reference individual (Derks et al., 2016).

## Results

### Approach summary

Our approach assumes two tissues, one pure tissue that serves as a reference and one mixed tissue of interest which is assumed to be “contaminated” with the pure tissue (Figure 1). The pure tissue can be easily isolated on its own, and the methylation levels and somatic mutations contained within it can therefore be easily determined by sequencing. The mixed tissue cannot be isolated on its own but the proportion of contamination can be determined using somatic mutations found in pure tissue allowing deconvolution of methylation levels.

**Figure 1:**
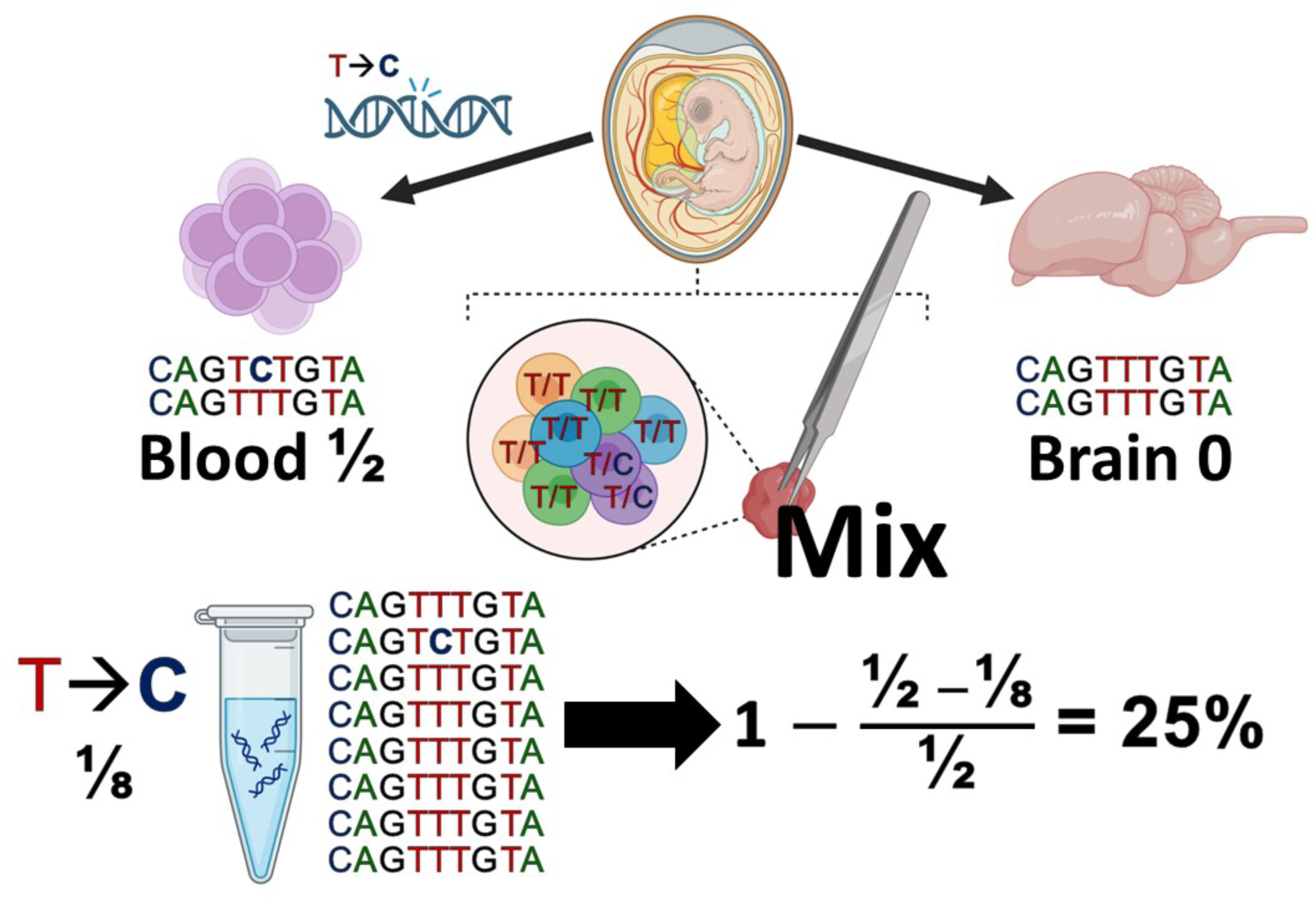
Illustration of the deconvolution approach based on novel somatic mutations. In this example, blood is the pure tissue and brain is a mixed tissue containing an unknown proportion of blood DNA. During the development of a bird a somatic mutation (T → C) occurs in hematopoietic (blood cell) progenitors but not the brain resulting in a blood-specific somatic-allele frequency of 50% (Top). Sampling brain tissue containing an unknown proportion of blood also samples the somatic mutation in proportion to the amount of blood cells included (Center). Extracting and sequencing this tissue allows for quantification of the germline and somatic mutation frequencies at this site revealing an allele frequency of ⅛ for the somatic mutation (Bottom). This provides for a 75% reduction in the frequency of the somatic mutation, revealing the tissue DNA to be 25% blood and 75% brain. Image in part with Created with BioRender.com.

Examples of applicable systems would include pure normal tissue intermixed with a tumor of interest, pure somatic tissue intermixed with germline tissue, or pure blood contaminating a highly-vascularized tissue of interest. To demonstrate this approach, we first confirm that somatic mutations in pure tissues should be ubiquitous and easily differentiated from germline heterozygous sites using tissue specific allele frequencies. We then show that changes in allele frequencies of somatic mutations can be used to predict the contributing portions of tissues in a mix. Finally, we show that the relative proportions of tissues in a mix can be used to deconvolute methylation states in CpG and non-CpG context and across genomic features.

### Somatic mutations are ubiquitous but at low frequency

Our approach relies on disentangling somatic mutations from germline heterozygous sites using differences in allele frequencies. To illustrate that our approach is in principle possible, we begin by characterizing the expected number and frequencies of somatic mutations in pure tissues with simulations. Somatic tissues are effectively populations of cells that expanded from smaller populations without recombination (De, 2011; Woodworth et al., 2017). We simulate the expected allele frequencies for somatic mutations using FastSimCoal2 (Excoffier et al., 2021). We simulate allele frequency information for populations of 100 somatic cells doubling every generation for 12 cycles of division. This reflects the the minimum possible number of divisions necessary to go from the 50,000 cells of a great tit epiblast (Nájera & Weijer, 2020) to the 226 million cells of a great tit brain (Olkowicz et al., 2016). Simulations were run across 11 mutation rates (µ), ranging from 2.3 x 10^-9^ mutations per base pair per division to 4.6 x 10^-8^ mutations per base pair per division, representing the range of germline mutation rates recorded across birds (Bergeron et al., 2023). Though reliable estimates of somatic mutations rates in birds are non-existent, somatic mutation rates in mammals are consistently higher than germline mutation rates indicating that our mutation rates should be very conservative (Cagan et al., 2022; Oota, 2020). Altogether, 100 simulations were run for each of the 11 values of µ and aggregated (N=1100) for downstream analyses with 50 diploid cells sampled for each simulation. The number of somatic mutations increased proportionally to µ (Poisson Regression: p<10^-6^, r^2^=0.791), ranging from as low as 578,808 somatic SNVs under a µ of 2.3x10^-9^ to as high as 38,504,756 somatic SNVs under a µ of 4.6x10^-8^ (Figure 2A). Allele frequency mean (Linear Regression: p=0.808, F=0.059, r^2^=0.000), median (Linear Regression: p=0.5178, F=0.4185, r^2^=0.000), and mode (Linear Regression: p=0.560, F=0.34, r^2^=0.000) were however unaffected by µ (Figure 2B). Somatic mutations were relatively rare across all simulations and mutations rates with mean frequencies per site of 0.182 (Range: 0.066-0.322), median values of 0.093 (Range: 0.02-0.49), and modal values of 0.0927 (Range: 0.1-0.92). We define low-frequency variants as those with minor allele frequencies (MAF) less than 0.25—a value arbitrarily chosen to be substantially less than the expected frequency of at germline heterozygous site. The vast majority (79.3%) of somatic mutations were low-frequency with a range of 35.7%-99.3% falling below the 0.25 MAF threshold across all simulations and mutation rates.

**Figure 2:**
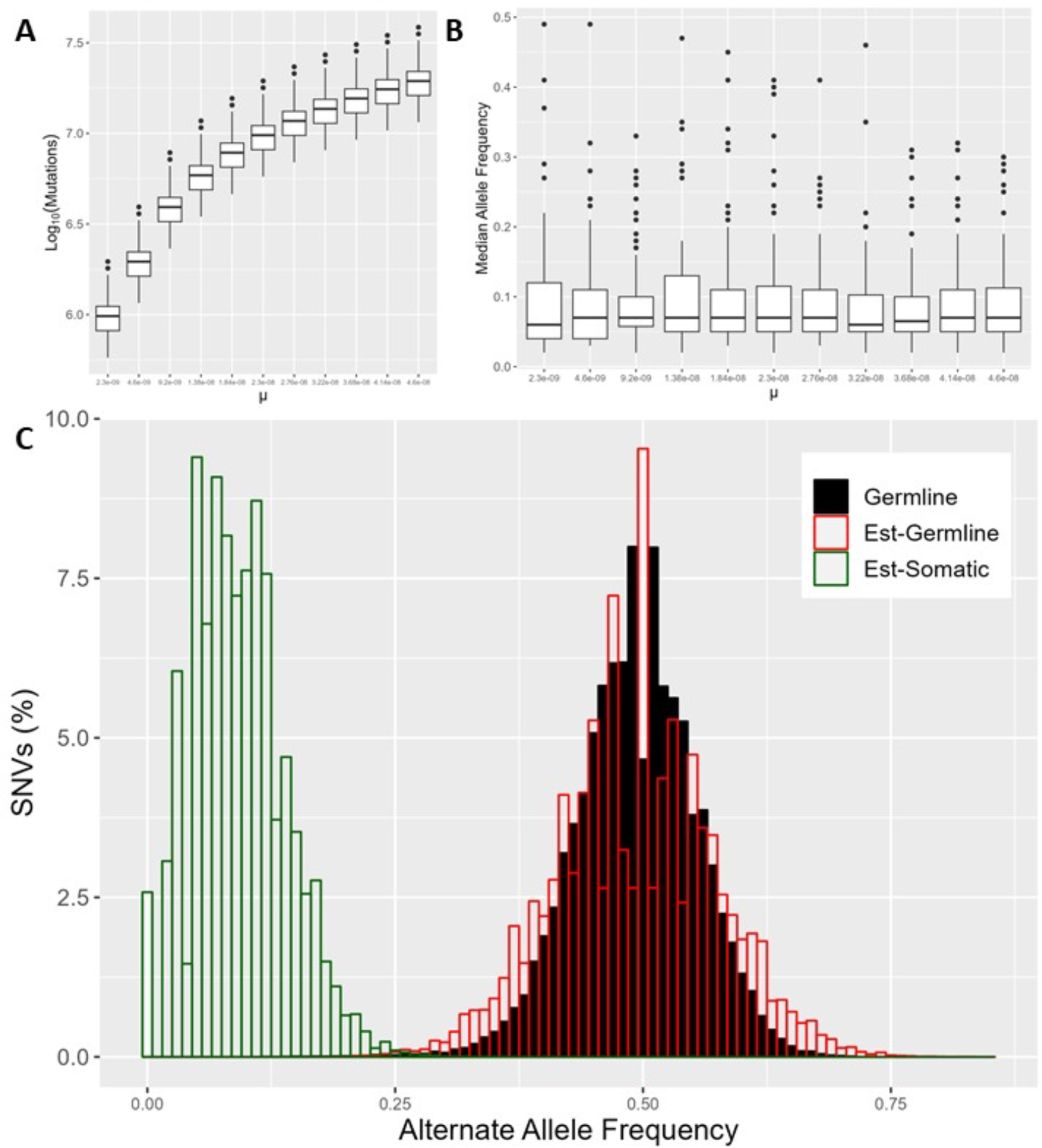
Distributions of novel somatic mutations and germline heterozygous sites sampled from a single tissue. **A) The number of novel somatic mutations scale with (somatic) mutation rate.** Shown are the numbers of observed somatic mutations in 100 simulation runs for varying mutation rates µ. **B) Median allele frequency is uninfluenced by mutation rate.** Boxplots indicate the median derived allele frequency across the 100 simulation runs made for varying mutation rate µ. **C) Overlap in allele frequency distributions of novel somatic and germline mutations is minimal.** Histogram of simulated germline heterozygous sites (Black, “Germline”) with expected distributions of germline (Red, “Est-Germline”) and somatic (Green, “Est-Somatic”) mutations at median frequency. Expected distributions are drawn from a binomial distribution at half the sampling frequencies of simulated sites, which provide a suitable estimate for occurrence of rare variants.

To confirm the expected distribution of germline heterozygous sites in sequencing data we simulated bisulfite short-reads from the great tit reference genome using the Sherman read simulation tool from the Babraham Institute (Gong et al., 2022; Grehl et al., 2020; Nunn et al., 2021). Germline mutations were introduced into the reference genome at a rate of one SNV every 1,000 sites using Mutation Simulator (Kuhl et al., 2021) prior to generating reads. Reads were generated at 400X in equal proportion for each haplotype, merged, and subsampled down to 100X before alignment back to the reference. We tested whether allele frequencies of simulated reads at germline heterozygous sites had a predictable binomial distribution, which could be used to differentiate them from somatic mutations. Germline heterozygous sites had a lower mean (0.493 versus 0.500) and median (0.494 versus 0.500), and were over-dispersed relative to a strict binomial distribution with higher standard deviation (0.081 versus 0.077) and absolute deviation from the median (0.076 versus 0.074) based on Monte Carlo tests (10,000 permutations, p<0.0001). Due to this, a higher proportion of sites contained low-frequency variants in the simulations relative (4.955x10^-3^) to a strict binomial distribution at the sampled depth (2.032x10^-3^). After filtering to remove sites that were in repetitive elements, segmental duplications, and more than twice or less than half of the average depth, the distribution of simulated germline heterozygous sites still showed the same slight downward shift in mean and median allele frequency relative to a strict binomial distribution. A higher proportion of sites remained at low-frequency in the simulations (1.609x10^-3^) relative to the binomial distribution (2.01x10^-5^). However, filtered sites showed a lower but similar dispersion to a binomial distribution taken at half depth with a comparable proportion of rare sites (Two-Sided Monte Carlo Test: p=0.9770; N=10,000). As such, a half-depth binomial model functions as a conservative null hypothesis for the distribution of germline heterozygous sites in bisulfite sequencing data. Comparison on these to simulated distributions of germline sites reveals minimal overlap with germline sites (Figure 2C).

### Great tit data: In silico mixtures of methylated blood and brain

We demonstrate and benchmark our approach using bisulfite-sequencing data obtained from blood and brain tissues of the great tit reference individual. As the reference individual was generated from an extensive library obtained from blood samples, we have the additional benefit of a detailed distribution of heterozygous sites of genomic DNA to verify our approach. In our example, blood is treated as the pure tissue and brain is treated as part of the “convoluted” mixed tissue (e.g. blood-brain mix). To verify and benchmark our approach we were interested in generating convoluted tissue from real sequencing reads for which we know the true level of contamination. To generate these mixed tissues, we first obtained all available raw reads, 292 and 358 million for blood and brain tissue, respectively. We then created eleven *in silico* subsamples of blood and brain reads ranging 0% blood (100% brain) to 100% blood (0% brain) in 10% increments in each tissue proportion (Figure 3A). Total read count was maintained at ∼292 million read pairs in each subsample. Total subsampled read count was effectively constant across all 11 of these groups. We then conducted a best-practice NGS workflow to obtain genomic along with DNA methylation variation.

**Figure 3:**
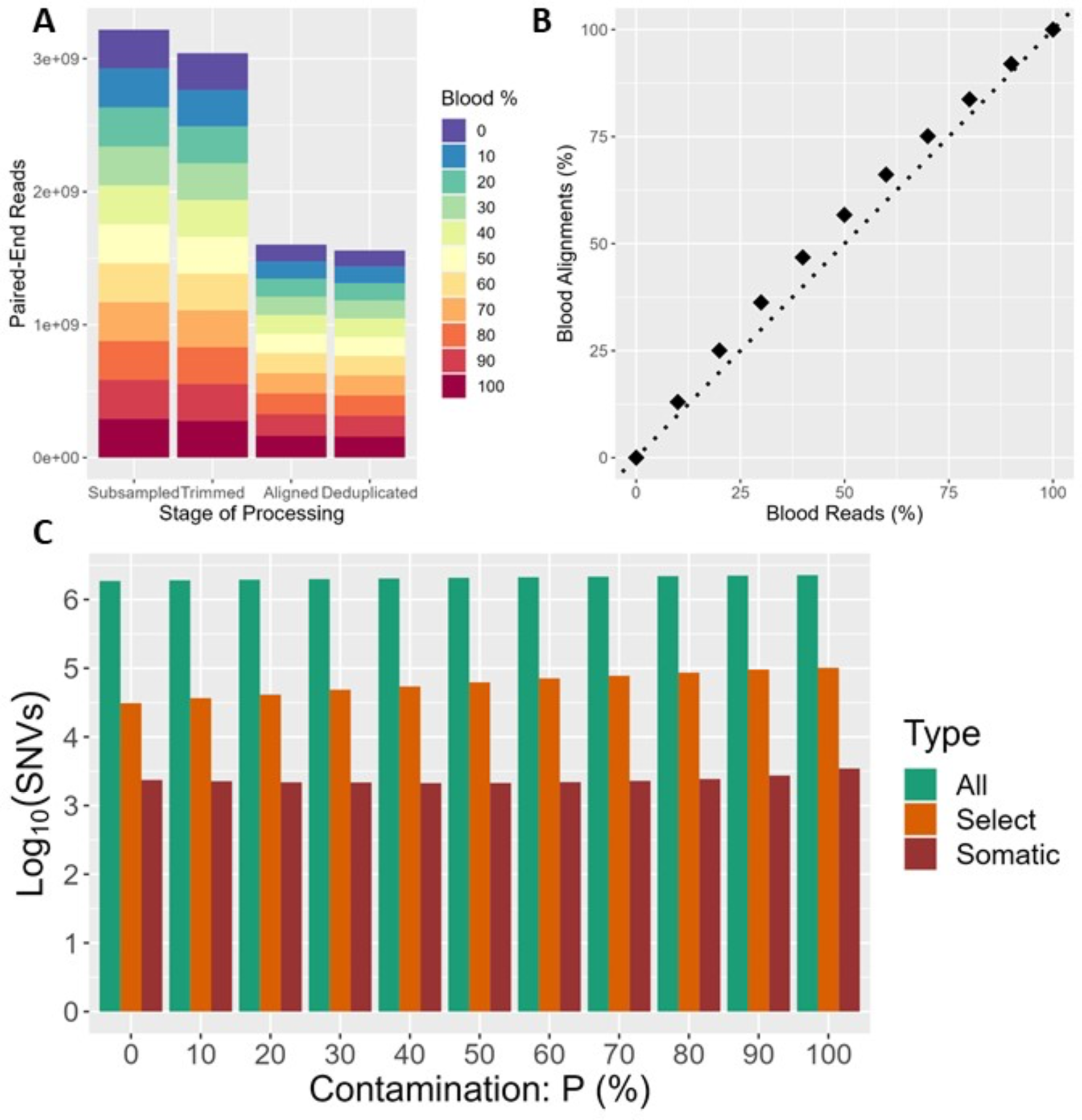
Validation of the “contamination” levels P of blood reads in blood-brain read mix. A) Stacked bar chart of the number of paired-end reads after each step of processing: subsampling, read trimming, alignment to the reference genome, and deduplication. B) Correspondence of final aligned and deduplicated reads on the (Y-Axis) by subsampled reads on the (X-Axis) with dashed line showing a perfect one-to-one relationship (r^2^=0.996). C) Grouped bar chart of Log_10_ number of SNVs for each *in silico* sampling by proportion of initial blood reads. Groups are color-coded by type of SNV: all SNV calls (All); select high-confidence SNV calls at high depth, and quality, with repeats filtered, and bisulfite-independent mutation types (Select); and SNVs at select sites classified as somatic in blood based on allele frequency (Somatic).

Preprocessing (Subsampling, trimming, aligning and deduplication) had very limited effect on the read data. Quality trimming had a minimal impact on differences in read count, but blood reads aligned to the reference genome more effectively producing a slight gradient from 253 million aligned reads in the uncontaminated pure brain to 329 million aligned reads in pure blood sample (Figure 3A). Because of this and slight variation in duplication proportions between mixes, the proportion of blood reads in the final alignment varied and was slightly higher than the original mixture proportions. Despite some difference in alignment efficiency between tissues subsampled, read proportions strongly predicted final aligned read proportions (Figure 3B; r^2^=0.994, p<10^-6^). Methylation percentage increased at CpG and non-CpG sites with the proportion of brain reads in each mixture ranging from 44.6% in pure blood to 49.4% in pure brain at CpG sites, and 0.1% in pure blood to 1.7% in pure brain at non-CpG sites, consistent with expectations of generally higher methylation levels in the brain (Derks et al., 2016).

### Somatic SNVs and proportion of contamination P

We called SNVs using BS-SNPer (Gao et al., 2015), which allows acquisition of sequence variation from bisulphite data. With this information we can differentiate and deconvolute mixtures using somatic SNVs. SNV calls varied between mixtures but increased proportionally to the percentage of blood reads included, ranging from 1,865,785 in pure brain to 2,267,659 in pure blood. This finding is driven by higher SNV counts in the blood reads than brain reads. Our deconvolution approach requires that we predict the relative proportion in which tissues are mixed using blood-specific somatic SNVs (Figure 3C). First, SNVs from pure blood were filtered to include only selected high-confidence sites outside of repetitive features. These high confidence sites were highly variable between mixtures and increased with the proportion blood reads, ranging from 31,018 sites in the pure brain “mix” to 101,459 sites in the pure blood “mix”. Somatic mutations were then identified at these selected sites based on allele frequency with an MAF below 0.25 in either the pure blood or the mixed tissues. We identified 2,123 to 3,465 somatic SNVs in blood across subsamples. These corresponded to approximately 0.0936%-0.153% of total blood SNVs. Comparison to heterozygous sites obtained from genomic reads (i.e. unconverted, non-bisulphite reads) at 90X from the same individual revealed that between 398 and 672 (18.5-19.5%) variant sites were found as low frequency in both data sets. Allele frequency of putative somatic mutations in blood reads also decreased proportionally as blood read percent dropped (p=<10^-6^; r^2^=0.987). This allowed for accurate prediction of the proportion of blood contamination P in each mixture with equation 1 (Figure 4). Our estimated P was strongly correlated with the real proportion blood reads in the initial subsamples (p= <10^-6^; r^2^= 0.985) and final alignment (p=<10^-6^; r^2^=0.994).

**Figure 4:**
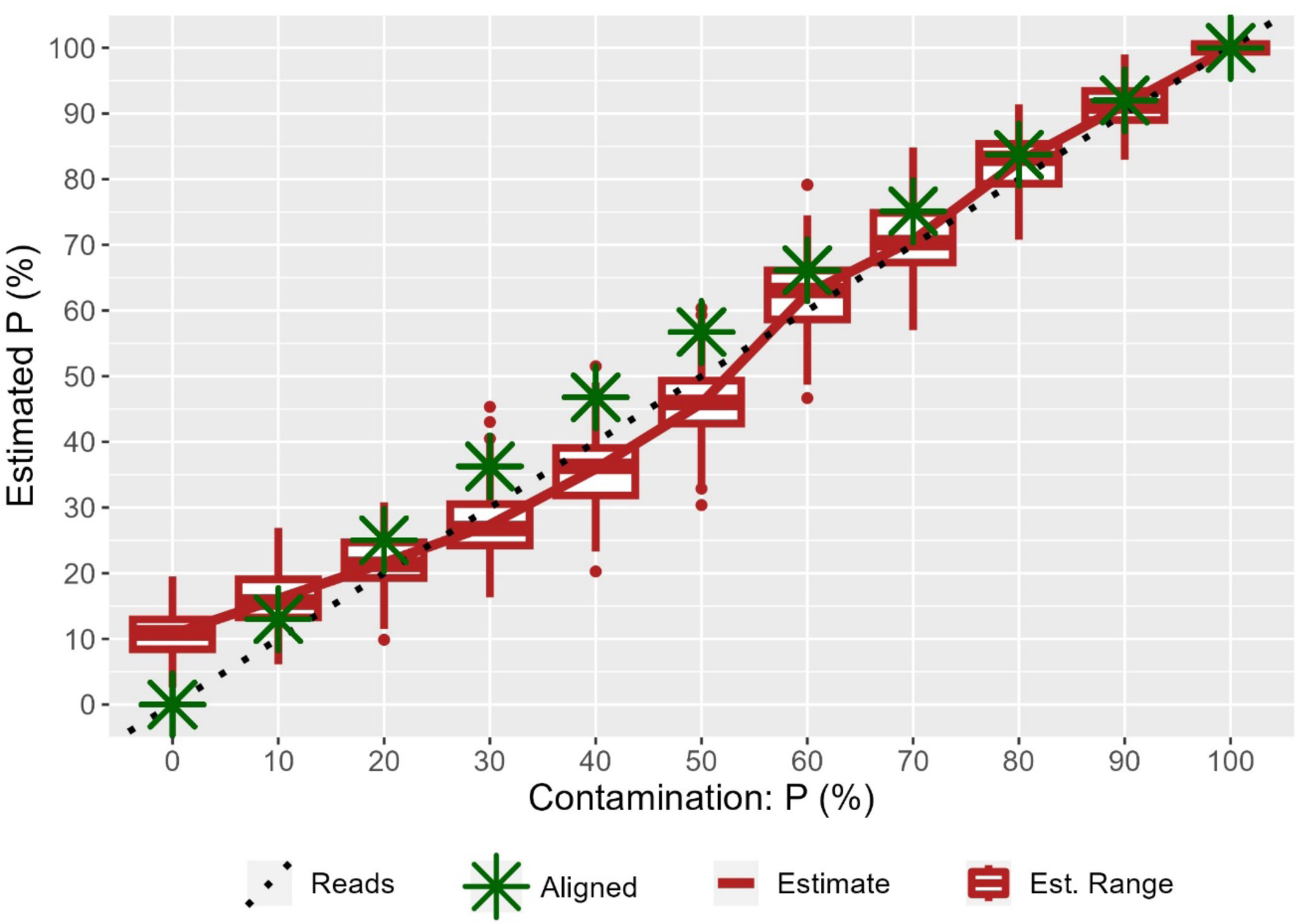
Estimating contamination proportion P using mutation frequencies in somatic tissue samples. Estimates of P based on the change in somatic mutation allele frequency. Proportion blood reads (“Reads”) in subsampled mixtures are shown with the dashed line, and proportion blood reads in the final deduplicated alignment (“Aligned”) are shown with green stars. The estimated proportion of blood reads (“Estimate”) is shown with a red line with a bootstrapped (n=100) range based on groups of 100 SNVs shown as boxplots (“Est. Range”).

### Deconvolution of methylation levels in contaminated tissues

After successfully recovering the level of contamination in the 11 mixed samples, we use the contamination proportion P to decompose DNA methylation levels from the mixed tissues. This method could be applied in principle if there would be any *a-priori* knowledge of P, e.g. obtained from a different method. However, we want to stress that tissue proportion may not reflect the relative proportion of genomic reads. We therefore highly recommend that P be obtained through *de-novo* mutations.

Principally we deconvolute methylation state across mixtures by regressing out pure tissue methylations at each site (Equation 2 and Equation 3) using the contamination proportion P obtained for a particular mix. We validate our approach through comparison of both convoluted and deconvoluted mixtures to pure brain estimates (Figure 5) using concordance correlation analysis (Lin, 1989). Validation at the site level used random subsampling of 30k cytosines, one from CpG and one from non-CpG context. The same sites were used for all subsamples to facilitate comparisons. As deconvolution is meaningless at non-methylated sites, only sites with methylation levels of at least 10% in pure blood or brain were sampled (Derks et al., 2016). Deconvolution resulted in significantly (Bootstrapping: Replicates=10,000; p<10^-4^) improved concordance correlation (ρ_c_) in CpG context for mixtures of 20-70% and in all mixtures for non-CpG context (Figure 5A). The magnitude of deviation from pure brain methylation state was likewise reduced for all mixtures regardless of context (Wilcoxon Signed Rank Tests: p<10^-6^, V> 48,581,408). Bias was estimated as Cb (=ρ_c_/ρ) the 0-1 scaled deviation of the slope from a perfect (1:1) correlation (Lin, 1989). Across sites, no bias (Figure 5B) was observed for deconvoluted estimates or convoluted mixture within CpG context. However, deconvolution resulted in substantially reduced bias in estimates of methylation state in non-CpG context (Figure 5B). Finally, deconvoluted reads were statistically concordant with pure brain reads at all deconvolution percentages, with >99% of sites possessing exact 99% confidence intervals that included the true methylation percent derived from pure brain reads, regardless of contamination percent or cytosine context (Figure 5C). In contrast, mixed methylation calls showed steadily decreasing statistical overlap with brain methylation calls as contamination percent increased with a more pronounced effect at CpG sites than non-CpG sites.

**Figure 5:**
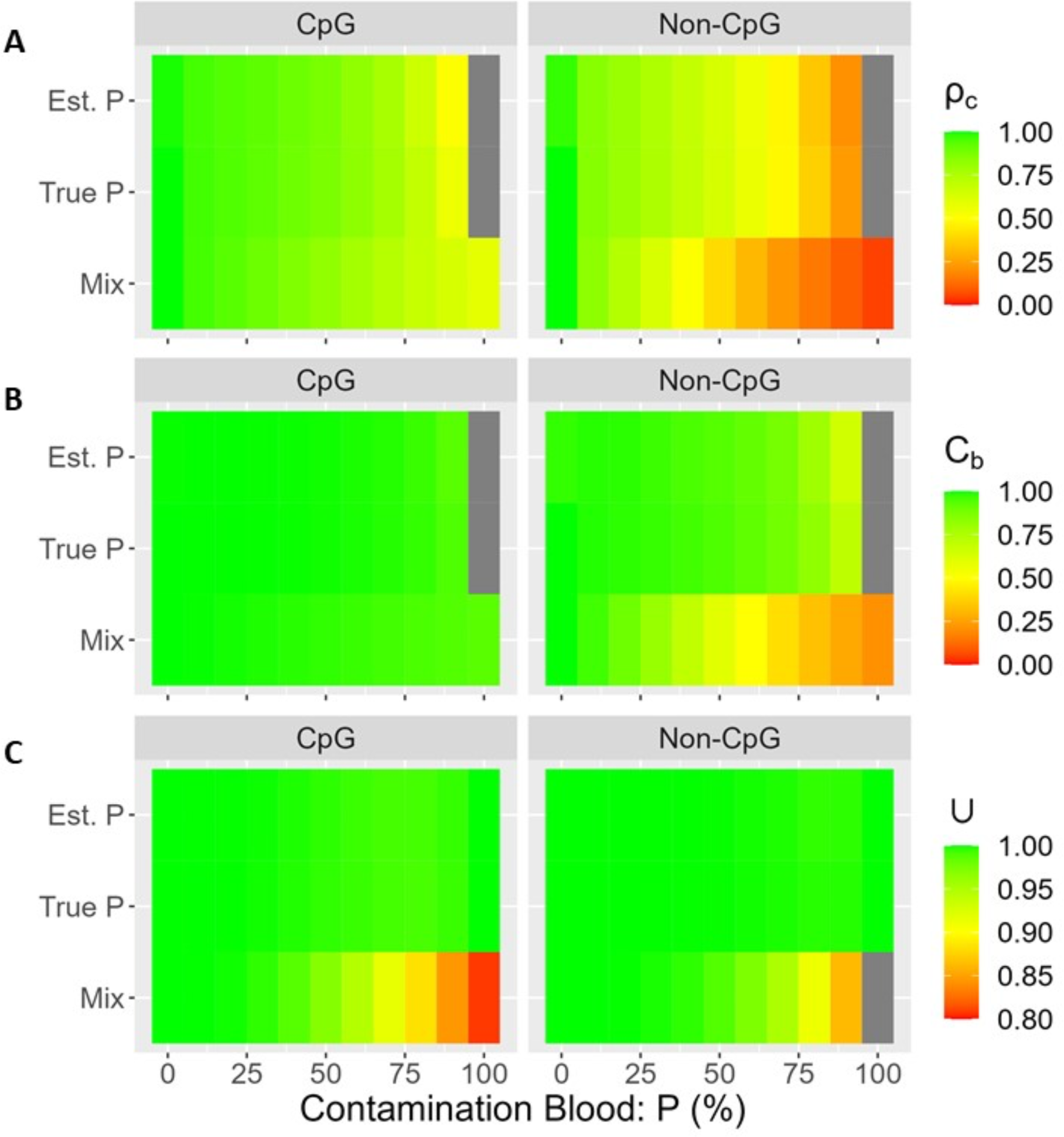
Deconvolution of site-specific methylation levels in mixed samples. A) Tile plots of correspondence correlation coefficients (Rho) of predicted versus actual methylation levels for deconvolutions and mixes in CpG (left) and non-CpG (right) context. B) Correspondence correlation C_b_ (a metric of bias direction in correlation representing deviation of origin, C_b_=ρ_c_/ρ) of predicted and actual methylation levels for deconvolutions and mixes in CpG (left) and non-CpG (right) cytosine context. C) Statistical concordance of predicted and actual methylation levels for deconvolutions and mixes in CpG (left) and non-CpG (right) cytosine context. Colors represent proportion of sites with methylation levels that are not statistically different from pure tissues at an alpha of 0.01.

We assessed the practical effectiveness of deconvolution in terms of its capacity to identify sites that were differentially methylated between blood and brain (Figure 6). Power was assessed as sensitivity (the proportion of true positives detected), and accuracy was assessed as specificity (the proportion of true negatives detected) using the pure blood and brain reads as a gold standard. These metrics were assessed at three types of features grouped by context—individual sites, whole-CpG islands, and whole gene bodies—and in both CpG and non-CpG context for six feature types total (Figure 6).

**Figure 6:**
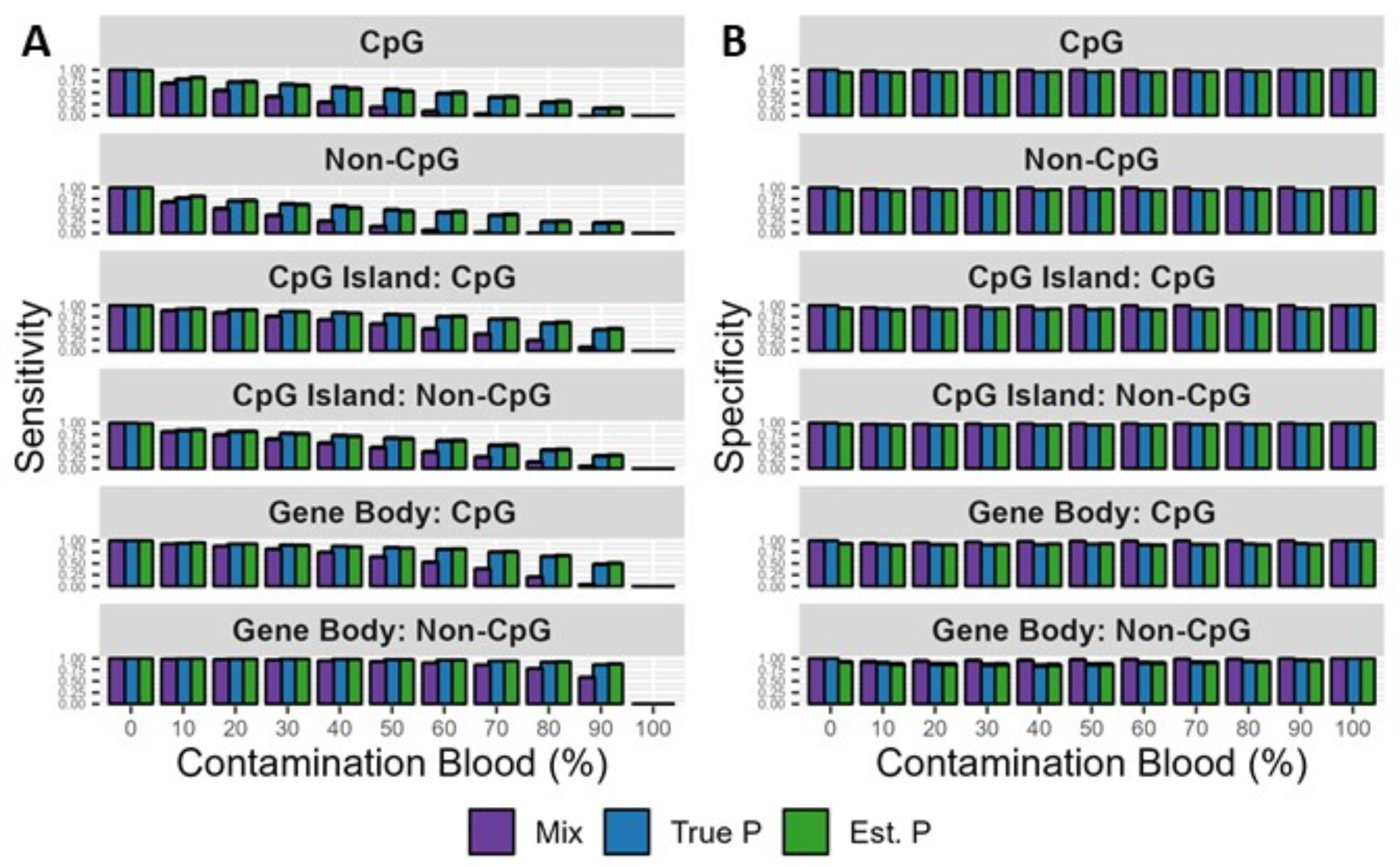
Sensitivity and specificity for methylation differences between blood and brain. Grouped bar chart of sensitivity (A) and specificity (B) for detection of methylated differences between the blood and brain tissue of great tits by deconvolution method (color), and context (facet) for cytosines above 10% methylation in either tissue. Sensitivity is defined as the proportion of true positive differences detected between the tissues. Specificity is defined as the proportion of true negative differences assigned to negative results.

Across all mixtures, features and context, (F=29.76; p<10^-6^; R^2^=0.8654), no significant interaction terms were observed between P, context, and impact of deconvolution suggesting that the effectiveness of deconvolution was similar across features and levels of contamination. Sensitivity varied by proportion contamination P, feature-type, and based on the application of deconvolution (F=124.6; p<10^-6^; R^2^=0.841). Sensitivity dropped precipitously as contamination levels increased (F=702.6; p<10^-6^; Partial-R^2^=0.609). This effect was seen for all feature types (F=45.21; p<10^-6^; Partial-R^2^=0.803) but recovery was generally better for larger features—excepting non-CpG methylation of CpG islands—than individual sites (Tukey Honest Significant Differences Post-Hoc: p<10^-6^). Deconvolution allowed for higher sensitivity regardless of feature type or cytosine context (F=33.89; p<10^-6^; Partial-R^2^=0.831)—however no difference was seen between estimates made based on true P and estimated P (Tukey Honest Significant Differences Post-Hoc: p=0.995) reinforcing that our method for estimating contamination levels with somatic mutations is highly accurate. At higher levels of contamination sensitivity was orders of magnitude higher with deconvolution than unmodified mixtures for all feature types. Identification of site-specific differences were effectively impossible (<10% detected) in convoluted mixtures above 50% regardless of context, but remained possible following deconvolution in all mixtures allowing identification of 17% of the truly different sites in CpG and 23% of these sites in non-CpG context even at 90% mixtures.

Regarding specificity, when comparing mixtures and features (F=9.12; p<10^-6^; R^2^=0.663), no significant interactions were seen between P, feature-type and the effects of deconvolution (Figure 6C). Significant differences were nonetheless observed based on the proportion of contamination, feature type and application of deconvolution (F=33.04; p<10^-6^; R^2^=0.583). Generally, specificity increased with increasing levels of contamination (F=23.03; p=3.22x10^-6^; Partial-R^2^=0.561)—likely due to lower depth and power (Gossmann & Waxman, 2022). Specificity differed by feature type (F=20.6; p<10^-6^; Partial-R^2^=0.465), with large features—excluding comparisons between non-CpG methylation and CpG islands—showing somewhat lower specificity than single sites (Tukey Honest Significant Differences Post-Hoc: p<10^-5^). Finally, use of raw mixtures rather than the deconvolution resulted in slightly higher specificity (F=70.52; p<10^-6^; Partial-R^2^=0.395). Deconvolutions based on true and estimated read counts did not differ in specificity (Tukey Honest Significant Differences Post-Hoc: p=0.238).

## Discussion

Our findings address the limitations of working with mixed tissues and demonstrate a straightforward and effective means to deconvolute methylation levels using tissue-specific somatic mutations. Use of mixed tissues biases the direction of methylation estimates, and prevents detection of differentially methylated sites. These biases are most pronounced at high levels of contamination and at single sites. Moreover, we note tissue specific variation in mapping success between tissue which might reflect differences in methylation pattern and biases of the reference genome—regardless of cause these introduce an additional complication into the use of mixed tissues. Applying our deconvolution approach to bisulfite sequencing data allowed for unbiased and accurate recovery of methylation states even at high levels of contaminating tissue in the mixture, and regardless of cytosine context. Our method greatly improved the sensitivity of detection of differentially methylated states at the levels of specific sites, CpG islands, and gene bodies. False positive rates across all sites and features were generally low and comparable to other methods for assessment of weakly methylated differences (Tran et al., 2018). Estimating methylation differences across CpG islands and gene bodies, resulted in higher false positive rates due to accumulation of error across sites. This problem was observed regardless of whether deconvolution was applied. Use of mixed tissues did however, result in generally lower false positive rates suggesting that methylation differences obtained with mixed tissues should still be trustworthy when detected. At higher levels of contamination use of mixed tissues is not a viable option, and deconvolution allows for recovery of most highly-differentiated sites and features with minimal error.

Our use of somatic mutations to assess levels of mixture between tissues allows for the deconvolution of tissues on generalized principles of mutation and inheritance. As such, methylation states can be deconvoluted without prior assumptions about patterns of epigenetic modification, making this method appropriate for use in non-model organisms. This contrasts with existing methods that rely on prior knowledge of cell-type specific methylation states (Loyfer et al., 2023; Moss et al., 2018; Zhu et al., 2022), or distributions of methylation patterns within an organism (Scherer et al., 2020; Schmidt et al., 2020; Zheng et al., 2014), which are generally lacking outside of humans and well-studied mammals (Aliaga et al., 2019; de Mendoza et al., 2020; Tanay & Sebé-Pedrós, 2021). Nor is our method restricted to analyzing canonical modifications at CpG sites. Indeed, emerging literature suggest that such canonical methylations may be of less overall importance than previously assumed (Derks et al., 2016; Fuso, 2018; Ramasamy et al., 2021; Yang et al., 2023), and our findings suggest that non-canonical methylations they may be more easily missed due to convolution and contamination of the focal tissue. Our method does require the use of a paired pure tissue alongside a contaminated tissue. It is appropriate to blood contamination which commonly occurs (Krausgruber et al., 2020; Liu et al., 2023; Vogel et al., 2019) and may be more pronounced in non-mammalian organisms with nucleated blood cells. Our method is also applicable to the study of tissues that are hard to isolate in pure form, such as tumors (Bergmann et al., 2016; Taylor-Weiner et al., 2018), germline tissues (Jenkins et al., 2017), embryonic tissues (Eckmann-Scholz et al., 2012; Morin et al., 2017), or those taken from smaller organisms across the eukaryotic tree of life (Lamka et al., 2022; Luo et al., 2018; Sturm et al., 2021).

As our method is not bound to methylation or epigenetic markers, it may conceivably be applied to the deconvolution of tissues for purposes other than the study of 5mC or similar epigenetic modifications (Talbert et al., 2019). Though tissue mosaicism (Lesaffre, 2021; Reusch et al., 2021; Rozhok & DeGregori, 2019) and epigenetic modification should be universal across multicellular organisms, 5mC modifications occur in only a limited (if diverse) subsampling or eukaryotes (Aliaga et al., 2019; de Mendoza et al., 2020; Schmitz et al., 2019).

Somatic mutations have been previously used to identify the relative purity and contamination of tumors and associated non-cancerous somatic tissues (Bergmann et al., 2016). Again, these approaches make assumptions about the selected tissues and the selected organisms. As we show, somatic mutations should be quite common and detectable in most tissues of most organisms—a conclusion that is all the more likely if somatic mutation-rates scale-negatively with life expectancy across organisms as has been proposed (Cagan et al., 2022). While previous efforts at somatic mutation detection have focused on high confidence calls at specific potentially important sites (Dou et al., 2018), our approach and simulations reveal the utility of somatic site-frequency spectra to highlight broad differences between tissues. In such context minor errors in the calling of somatic mutations are unimportant, and tissues can be studied in population genomic context.

## Conclusion

Ultimately, tissue heterogeneities constitute a neglected evolutionary interface in which genomic differences manifest as (potentially-selected) phenotypes. Evolutionary biology seeks to understand generalized patterns that persists across all or most organisms. Current methods and foci for research on tissue heterogeneity are however limited to select-few organisms, unsuitable for general applications, and constrained against application to evolutionary biology. By linking patterns of tissue mosaicism to universal patterns of mutation and inheritance, we may allow for expanded studies on tissue heterogeneity in the fields of epigenetics and evolutionary biology.

## Methods

### Simulations

Simulations of allele frequencies for somatic mutations were run in FastSimCoal2 (Excoffier et al., 2021). Simulations ran over 12 generations of divisions, in accordance with the minimum number of potential cell divisions from a great tit epiblast of 50,000 cells (Nájera & Weijer, 2020) into a great tit brain of 226 million cells (Olkowicz et al., 2016). Initial population size was set to 100 diploid cells (200 haploid genomes) which doubled every generation based on a parametrized continuous backward population growth rate of -0.69 (e^-0.69^∼0.5). While reliable estimates for somatic mutation rates for birds are lacking, mutations were set to vary one order of magnitude across the spectrum of germline mutation rates reported for birds (Bergeron et al., 2023). Studies on mammals indicate that somatic mutation rates are often orders of magnitude higher than germline mutation rates (Cagan et al., 2022; Oota, 2020), so our values are likely conservative estimates. Final sampling of somatic sites in the simulated brain was set to 100 haplotypes from 50 diploid cells.

Simulations of null heterozygous frequencies were run using Mutation Simulator (Kuhl et al., 2021) and the Sherman read simulation tool from the Babraham Institute (Gong et al., 2022; Grehl et al., 2020; Nunn et al., 2021). The reference genome from the great tit ‘Abel’ was downloaded from RefSeq and mutations were introduced such that an SNV was introduced at approximately 1 in every 1000 sites and insertions and deletions of up to 8 base pairs were each introduced every 1 in 20,000 sites. Sherman was then used to simulate 1,020,310,770 bisulfite read-pairs from both the reference genome and the simulated haplotype of this genome produced with Mutation Simulator. Sherman was run in paired-end mode with an error rate of 0.01% an insert size of 200-600 base pairs, a read length of 100 base pairs, a cytosine conversion of 54% in CpG context and 99% in non-CpG context. Reads produced with the mutated and original haplotypes were then merged and randomly subsampled from ∼400X to ∼100X depth using SeqTK with a seed of 23. Simulated reads were then processed in a manner identical as applied real data below.

### Data, Methylation, and Base Calling

The great tit reference genome and reads were downloaded from RefSeq (Accession: GCF_001522545.3) and Sequence Read Archive (SRA)— Accessions: SRR2070790 (Brain) and SRR2070791 (Blood)—respectively (Derks et al., 2016). Reads from SRA were downloaded and extracted using the SRAtoolkit suite (Leinonen et al., 2011). These consisted of 292,381,018 and 358,255,601 paired-end bisulphite converted reads taken respectively from the blood and brain tissues. Blood and brain reads were randomly subsampled with Seqtk (Shen et al., 2016) and mixed into proportions ranging from 0 to 100% of the respective tissues in 10% intervals. Mixed Fastq files were trimmed for removal of Illumina adapters and quality using Trimmomatic (Bolger et al., 2014): adapters were removed in palindromic search mode with two mismatches allowed, a minimum alignment score of 26 for palindrome clips and 8 for simple clips, and a minimum adaptor length of 4 basepairs; quality trimming removed trailing and leading bases with Phred quality scores of less than 3.

Quality trimmed reads were aligned to a bisulfite-converted reference genome (Abel) using Bismark to produce BAM files (Krueger & Andrews, 2011). Bismark alignments were performed using Bowtie2 (Langmead & Salzberg, 2012) and “end-2-end” alignment mode with a minimum seed length of 32 base pairs. BAM files were deduplicated using the “deduplicate_bismark” utility included with Bismark. Deduplicated BAM files were sorted with Samtools and used for input into BS-SNPer (Gao et al., 2015) for base-calling with the following calling options: minimum coverage of 1, maximum coverage 100, minimum mutation coverage of 1, mapping quality 20, maximum error rate 0.02, minimum heterozygous frequency of 0.01, and minimum homozygous frequency of 0.85. Unsorted BAM files were used for input into the “bismark_methylation_extractor”, run in pair-end mode with overlap between paired-end reads included in calls, and minimum coverage (i.e. “cutoff”) of 1. The first and last 2 base pairs of each read were excluded from calls to prevent bias associated with bisulfite conversion library preparation.

### Somatic SNVs

The proportion of reads attributable to blood in each mixture was assessed through estimated coverage at blood-associated somatic mutations. High-confidence candidate sites were first identified in pure blood. SNV calls from pure blood were filtered to remove repetitive elements, segmental duplications, reads with Phred quality scores below 30, depth below half or above twice the mixture average, CpG sites, and multiallelic sites. BS-SNPer calls SNVs using different coverages on Watson and Crick strands as a way to disentangle real SNVs from bisulphite converted SNVs (Gao et al., 2015). To avoid downstream problems resulting from these coverage estimates all SNVs called with 1-fold difference in depth between Watson and Crick strands were removed, as were all SNV types that require differential handling of Watson and Crick strands following bisulfite conversion, leaving only four mutation types: A→T, T→A, C→A, and G→T. Finally, to avoid regions of high mutation rates, poor mapping, and potentially redundant somatic mutations, we filtered all SNVs within 100,000 base pairs of another SNV.

Somatic mutations unique to blood were identified using allele frequency data. Filtered sites were further refined to remove any SNVs calls that occurred in either the mixed tissue or blood tissue at an alternate allele frequency outside the range of 0.25 to 0.75. This range was chosen based on probability of a site being non-germline heterozygous.

Proportion P was estimated for each mix by measuring the proportional drop in allele frequency of remaining (putative) somatic mutation sites between the mixture and the pure tissue.

### Deconvolution with proportion of contamination P

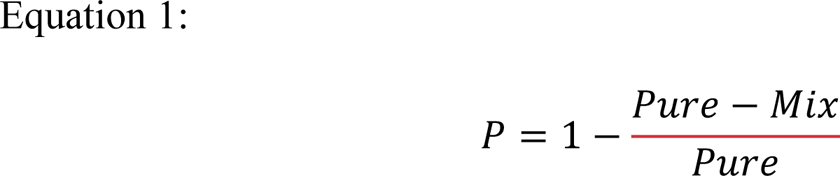

*P=*Proportion of contamination of pure tissue in mixed tissue

*Pure=*Minor Allele frequency at sites of pure-tissue somatic mutations in pure tissue

*Mix=*Minor allele frequency at sites of pure-tissue somatic mutations in mixed tissue

Deconvolution was performed by subtracting the proportion contaminating reads at each site in the mix tissue in proportion to the relative methylated and unmethylated reads at that site in the pure contaminating blood tissue (Equation 2 and 3). These methods were applied using the estimated P and the true P based on read count. Negative final values were changed to zero.

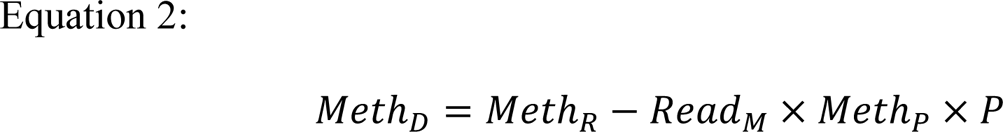

*Meth_D_*=Deconvoluted methylated reads

*Meth_R_*=Methylated reads in mixed tissue

*Reads_M_*=Total reads in mixed tissue

*Meth_P_*=Proportion of methylated reads in the mixed tissue

*P*=Proportion contamination of pure tissue in mixed tissue

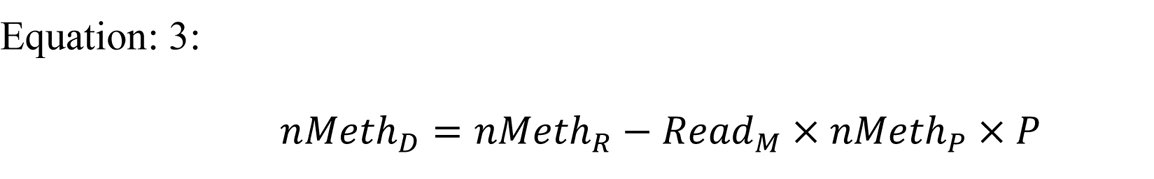

*nMeth_D_*=Deconvoluted non-methylated reads

*nMeth_R_*=Non-methylated reads in mixed tissue

*Reads_M_*=Total reads in mixed tissue

*nMeth_P_*=Proportion of non-methylated reads in the mixed tissue

*P*=Proportion contamination of pure tissue in mixed tissue

### Data Processing and Statistics

Samtools, Bedtools, and GNU tools suites were used for the general processing of genomic data (Danecek et al., 2021). Calculations of the proportion contamination P and deconvolution were performed with GNU tools AWK. General statistical analyses and preparation of figures were performed in R (R Core Team, 2023) using the RStudio Build 494 (Posit, Boston MA, USA) console on Windows 10. The following packages were used: Repeats were annotated using RepeatMasker run with HMMER and the DFAM3.7 database (Hubley et al., 2016; Tarailo-Graovac & Chen, 2009). Segmental duplications were annotated using BISER (Iseric et al., 2022). CpG island annotations were made with EMBOSS (Rice et al., 2000) and existing gene body annotations were used after downloading from RefSeq (Derks et al., 2016). All independent statistical hypothesis tests were conducted using a Dunn-Šidák corrected α of 0.00126 based on an initial α of 0.01 and eight independent tests (Somatic Mutations and Mu, Somatic Mutations and Distributions, Distribution of heterozygous sites, Read-x-Alignment Corr, P-x-Reads, Methylation by sites correlations, Sensitivity, Specificity). P-values for pairwise or grouped statistical hypotheses tests were calculated using a Bonferroni Correction when necessary. Pipelines for data processing, and statistics are provided on Github and in the supplementary materials.

## Data and resource availability

No new data were generated for this work. All code and script for analysis are available at https://github.com/JustinJonSchaderWilcox/Deconvolution or as supplementary materials.

## Acknowledgements

This project has received funding from the European Research Council (ERC) under the European Union’s Horizon 2020 research and innovation programme grant agreement No. 947636. This work was performed using the Linux HPC cluster at Technical University Dortmund (LiDO3), partially funded in the course of the Large-Scale Equipment Initiative by the Deutsche Forschungsgemeinschaft (DFG, German Research Foundation) as project 271512359. The author’s gratefully thank the LiDO3 team. This work was also performed with the support of the BMBF-funded de.NBI Cloud within the German Network for Bioinformatics Infrastructure (de.NBI) (031A532B, 031A533A, 031A533B, 031A534A, 031A535A, 031A537A, 031A537B, 031A537C, 031A537D, 031A538A).

## Author contribution

TIG designed and supervised the study. JJSW designed the simulations and conducted the majority of the analyses. QJRF conducted preliminary analyses. JJSW drafted the manuscript with input from all of the authors. JJSW and TIG finalized the manuscript with feedback from QJRF.

